# Proteomic Analysis of Human Chronic Traumatic Encephalopathy Brain Implicates Proteasome and Ribosome Dysfunction in Disease Progression

**DOI:** 10.64898/2026.02.13.705123

**Authors:** Helen E. Pennington, Dillon Shapiro, Jenny Empawi, Nurgul Aytan, Victor Alvarez, Jessie Mez, Michael L. Alosco, Xiaoling Zhang, Ann C. McKee, Thor D. Stein, Jonathan D. Cherry, Adam Labadorf

## Abstract

Chronic traumatic encephalopathy (CTE) is a progressive neurodegenerative disease associated with repeated head injuries (RHI) commonly experienced by contact sport athletes, military personnel, and domestic abuse victims. Despite growing recognition of CTE, the molecular mechanisms underlying disease progression remain poorly understood. This study aims to identify proteomic alterations associated with CTE pathology and clinical features to elucidate key biological pathways involved in disease pathogenesis. SomaScan 7k high-throughput proteomics was performed on 204 dorsolateral prefrontal cortex samples from the Boston University CTE Center Brain Bank. We identified differentially expressed proteins associated with CTE, hyperphosphorylated tau (ptau) pathology, duration of contact sports play, dementia status, and Cognitive Difficulty Scale (CDS) scores. Gene set enrichment analysis revealed that proteasome subunit proteins and related pathways were strongly associated with CTE progression and correlated with years of contact sports play. Reduction in ribosomal proteins and pathways was closely associated with ptau burden. Additionally, multiple models demonstrated significant alterations in MAPK-related cell signaling pathways. These findings advance our understanding of CTE progression and identify mechanisms correlated with key pathological features of the disease. Validation of these results could inform the development of diagnostics and treatments for CTE.

## Introduction

Chronic traumatic encephalopathy (CTE) is a progressive neurodegenerative disease associated with exposure to repeated head impacts (RHI).^1^ Millions of people experience RHI exposure every year through contact sports, military service, or domestic abuse.^4^ The primary risk factor for developing CTE is the cumulative force of RHI over an individual’s life, which varies depending on force, frequency, and duration of head impacts.^1,5^ Pathologically, CTE is classified as a tauopathy and characterized by the perivascular accumulation of hyperphosphorylated tau (ptau) in neurons.^1^ Abnormal ptau first appears at the depths of the cortical sulci and later spreads to other brain regions.^1^ Currently, CTE can only be definitively diagnosed through postmortem autopsy. The stage of CTE is determined by assessment of the distribution of tau tangles across the brain, and the degree of ptau aggregation is directly associated with the dose and duration of RHI exposure.^6^ Clinically, CTE presents with cognitive and behavioral impairments as well as mood disturbances and motor dysfunction.^2^

Much remains unknown about the molecular mechanisms that underlie CTE. Previous work studying transcriptional changes in human brain revealed that increased neurodegeneration, neuroinflammation, and decreased expression of certain synaptic markers were associated with disease progression.^3,7^ Recent targeted proteomic studies evaluating protein changes in CTE reported similar findings, also noting evidence of neurodegeneration, inflammation, and glial proliferation that bore similarities with Alzheimer’s disease (AD) pathology.^8–10^ These studies identified additional changes associated with CTE pathology, including a decrease in axonal guidance proteins, dysregulation of cell signaling proteins, and disruption to vascular and extracellular matrix (ECM) structures at certain disease stages.^8–10^ However, these studies were limited in sample size and scope, motivating further investigation of unbiased proteomic data in a larger cohort of individuals in relation to key clinical, exposure, and pathological measures of disease.

In this study, we present a bioinformatic analysis of SomaScan 7k high-throughput proteomics data from 204 prefrontal cortical brain tissue samples donated by players of contact sports to the Understanding Neurologic Injury and Traumatic Encephalopathy (UNITE) BU CTE Center brain bank. The goal of the study was to characterize differential protein abundance associated with important clinical factors, including cortical tau pathology burden, duration of play, CTE severity as assessed using the McKee staging criteria,^11,12^ dementia status, and Cognitive Difficulty Scale (CDS) score. Gene set enrichment analysis of significantly associated proteins provided a broader view of the biological processes implicated by the differentially expressed proteins. An integrated network analysis of the results provided insight into the interrelationships among the clinical, exposure, and pathological characteristics that were considered. Together, these analyses aim to shed light on the proteomic changes that underly CTE pathology and symptoms and suggest diagnostic and therapeutic targets for further study.

## Methods

### Brain donors

Brain donors with a history of repetitive head impact or traumatic brain injury exposure, and with frozen tissue from the dorsolateral prefrontal cortex available (n=248) were drawn from the Understanding Neurologic Injury and Traumatic Encephalopathy (UNITE) study. Inclusion criteria for UNITE included a history of contact sports participation, military service, or domestic violence and have been described previously, and further filtered by criteria used in our previous RNAseq study.^7,18^ Established criteria were used to diagnose Alzheimer’s disease (NIA-Reagan), limbic/neocortical Lewy body disease (LBD), and neuropathological presence of frontotemporal dementia (FTD).^13–15^ CTE was neuropathologically diagnosed using the NIH consensus criteria, which include abnormal perivascular accumulations of hyperphosphorylated tau (p-tau) in neurons, with or without astrocytes, and cell processes concentrated at the sulcal depths.^12^ Staging was based on regional p-tau involvement according to the McKee staging system since this has been shown to correlate with duration of play and clinical symptoms.^11,12^ Neuropathologists provided all disease evaluations while blinded to clinical evaluations. Brain donors were divided into three main groups: RHI controls (cases with a history of RHI but without a diagnosis of CTE) (*n* = 58), stages I and II (low CTE) (*n* = 33), and stages III and IV (high CTE) (*n* = 113). Demographic details, incidence and type of comorbid pathology, and contact sport histories are presented in Table 1.

**Table 1.** Demographic and Clinical Characteristics. Data are presented as mean (SEM) for continuous variables and % (*n*) for categorical variables. ANOVA was used to assess *P*-values for continuous variables, and chi-squared tests were used for categorical variables.

| Characteristic | RHI<br>($n = 58$) | Low CTE<br>($n = 33$) | High CTE<br>($n = 113$) | All Cases<br>($n = 204$) | *p*-value |
| --- | --- | --- | --- | --- | --- |
| **Cohort** |  |  |  |  |  |
| **Age at death, years** | 65.6 (2.4) | 56.0 (3.12) | 75.6 (0.97) | 69.7 (1.1) | **<0.001\*** |
| Range | 18–99 | 25–84 | 32–98 | 18–99 |  |
| **Post-Mortem Interval (PMI), hours** | 36.9 (3.22) | 45.3 (5.46) | 31.9 (2.00) | 35.4 (1.71) | **0.0217\*** |
| **Duration of contact sports play, years** | 10.6 (1.0) | 11.88 (0.96) | 16.2 (0.62) | 14.2 (0.50) | **<0.001\*** |
| American Football | 67.8% (40) | 93.8% (30) | 89.6% (103) | 84.4% (173) |  |
| Ice Hockey | 0.0% (0) | 0.0% (0) | 2.6% (3) | 1.5% (3) |  |
| Wrestling | 1.7% (1) | 3.4% (2) | 0.9% (1) | 2.0% (4) |  |
| Boxing | 6.8% (4) | 0.0% (0) | 2.6% (3) | 3.4% (7) |  |
| Soccer | 1.7% (1) | 0.0% (0) | 2.6% (3) | 2.4% (5) |  |
| Other | 20.3% (12) | 0.0% (0) | 1.7% (2) | 6.8% (14) |  |
| Alzheimer's Disease (AD) | 25.4% (15) | 3.1% (1) | 32.2% (37) | 25.9% (53) | **0.017\*** |
| Frontotemporal Dementia (FTD) | 8.6% (5) | 0.0% (0) | 10.4% (12) | 8.8% (18) | $\square 0.205$ |
| Lewy Body Dementia (LBD) | 19.0% (11) | 6.3% (2) | 24.4% (28) | 20.5% (42) | $\square 0.131$ |

### Tau Pathology

An AT8 antibody (Invitrogen MN1020, 1:1000) was used for immunostaining ptau as previously described.^16^ Ptau levels in the dorsolateral frontal cortex were determined based on AT8+ cell density analysis performed on formalin fixed, paraffin embedded tissue blocks harvested from the contralateral hemisphere brain region used for proteomics. The HALO AI v4.0 digital histology platform was used to quantify the amount of total ptau in the samples using standard methods described by Kanner et al.^17^ These AT8 total values were then normalized by taking the natural logarithm and adding the smallest AT8 total value.

### Clinical Details

Clinical information and sports history were evaluated based on interviews and questionnaires completed by the next of kin as previously described by Mez et al.^18,19^ Phone interviews were utilized to determine sports played and duration of play. Dementia diagnoses were determined based on modified DSM-IV criteria.^20^ The Cognitive Difficulty Scale score (CDS) evaluates cognitive impairment, with higher scores indicating greater impairment.^21^ The total score (CDS) was used in all relevant models.

### Covariate Imputation

Single imputation was performed using the Amelia R package^22^ to substitute 24 missing Postmortem Interval (PMI) values and one AD status value. All available covariates with fewer than 30 missing cases for that feature were used to impute the missing PMI and AD values using an empirical prior of 0.005 multiplied by the number of samples (204). Imputed covariate values were used for statistical adjustment in all models. Primary model variables (i.e., CTE status, duration of play, AT8, CDS, and dementia status) were not imputed (Table 2).

**Table 2.** Summary of Proteomics Findings. The table outlines the number of samples for each of the six main models as well as the number of significant proteins and pathways identified for each model.

| Model | Samples included | Proteins $p < 0.05$ | Pathways $p < 0.05$ | Pathways FDR $< 0.05$ |
| --- | --- | --- | --- | --- |
| AT8 Total | 180 | 109 | 228 | 30 |
| Duration of play | 194 | 588 | 230 | 141 |
| RHI to Low CTE | 91 | 58 | 391 | 192 |
| RHI to High CTE | 171 | 262 | 450 | 210 |
| Cognitive Difficulty Score | 132 | 154 | 282 | 37 |
| Dementia | 204 | 1008 | 455 | 215 |

### SomaScan 7k Proteomics

Approximately 20 milligrams of frozen tissue from the gyrus of the dorsolateral frontal cortex were collected from all 204 brain donors. The dorsolateral frontal cortex was selected because it is one of the brain regions where ptau develops first in the progression of CTE. Brain tissue from all 204 brain donors was lysed using a TissueLyser II instrument (Qiagen) in ice-cold Tissue Protein Extraction Reagent buffer (TPER, Thermo catalog #78510) with 1x Halt protease inhibitor cocktail (Thermo catalog #87786). One stainless steel bead (5 mm, Qiagen catalog #69989) was added to each round bottom sample tube (2 mL, Qiagen catalog #990381) containing a single tissue and TPER buffer plus protease inhibitor. Protein concentration of soluble protein was determined using the Pierce BCA protein assay (Pierce, Thermo catalog #23225). Equal amounts of total protein (2.4 μg per sample) were run in the cell and tissue lysate v4.1 SomaScan assay kit following the standard protocol recommended by the manufacturer (SomaLogic, Inc., Boulder, CO, USA).^23^ The final data were calibrated and normalized by SomaLogic Bioinformatics using their standard protocol and delivered as .adat files. All aptamers were maintained for every protein used in this study.

### Filtering and Preprocessing

The initial SomaScan 7k output contained data for 248 samples and 7,596 protein aptamers prior to quality control exclusions. First, calibrators, buffers, pooled samples, and two additional samples that were not present in the metadata were removed, leaving 210 brain donors. Next, all count data were log2-transformed, and four outlier samples were identified with standard deviations greater than 3 using Principal Component Analysis (PCA) PC1 as shown in Supplementary Fig. 1. These outliers were removed from the analysis. One female sample and one sample with incomplete metadata were excluded, resulting in 204 final study samples. After removing protein analytes labeled as non-human and “Internal Use Only,” 7,285 protein aptamers remained for analysis. Because the amount of clinical background information available varied among cases, the number of cases included in each model varied and is recorded in Table 2. Finally, the filtered data were converted from the .adat format to a summarized experiment format for use in RStudio. Sample and analyte filtering of the SomaScan 7k output is depicted in Fig. 1.

**Figure 1.**
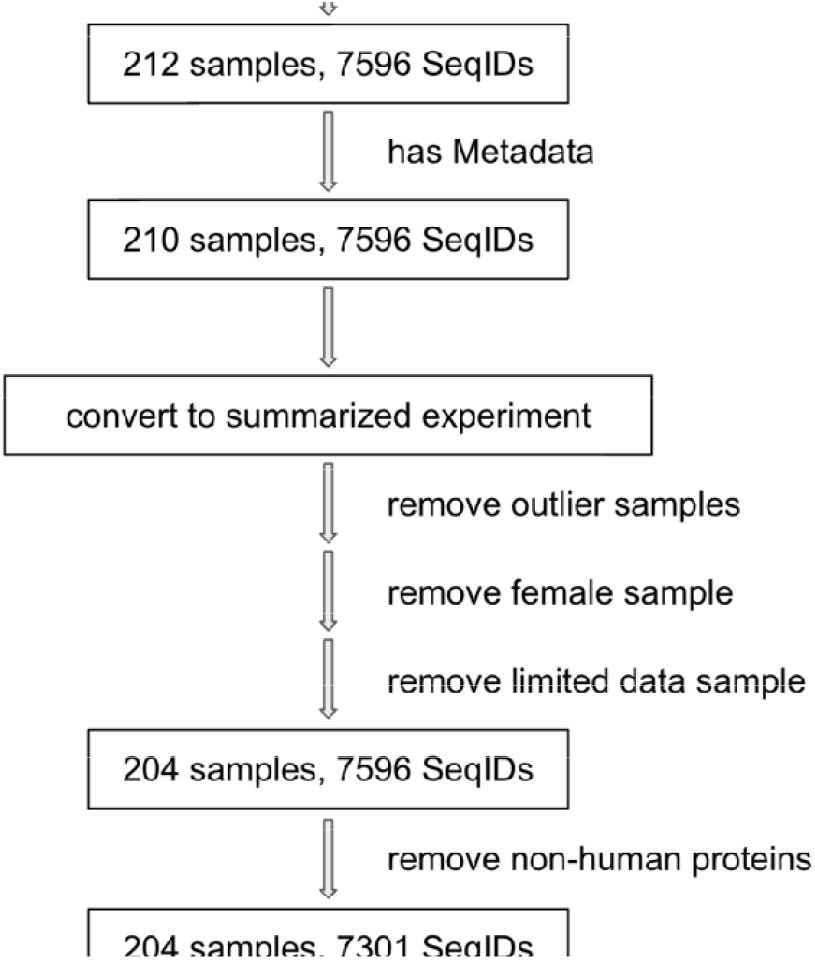
SomaScan 7k Proteomics Filtering and Preprocessing. Flow chart of SomaScan 7k proteomics filtering. Boxes represent updated numbers of samples and proteins included in the study. Arrows detail the filtering and transformation steps taken to arrive at the final dataset used in the study.

### Differential Protein Expression

Differential protein expression was conducted using the Limma software package.^23–25^ Separate Limma analyses were conducted for six main models: RHI vs. low CTE, RHI vs. high CTE, AT8 total, duration of play, CDS, and dementia status. Age of death, PMI, AD status, LBD status, and FTLD status were included as covariates for all models.^26^ Individual proteins were considered significant at a nominal *P*-value less than 0.05, as no proteins reached significance after Benjamini–Hochberg multiple testing correction in any model.

### Gene Set Enrichment Analysis

Gene set enrichment analysis (GSEA) was conducted using the MSigDB C2 Canonical Pathways gene sets for each model. Specifically, proteins were ranked based on descending log fold change in each Limma model, and the fgsea package was used to perform all GSEA analyses.^27,28^ All gene set enrichment analyses were subject to Benjamini–Hochberg (BH) multiple hypothesis adjustment, and gene sets with adjusted *P*-value less than 0.05 were considered statistically significant.

### Hierarchical Clustering

Statistically significant pathways were grouped from all six models into functional categories based on hierarchical clustering of their leading-edge proteins using the cosine similarity score method. The pathways were then sorted into larger-order groups using the cutreeDynamic function in the dynamicTreeCut R package and further refined by manual inspection based on gene content of the leading-edge genes. Groups were required to contain at least five pathways (Supplementary Fig. 4). A clustered heatmap depicting the hierarchical clustering is also included in Supplementary Fig. 4. Groups were assessed and named based on their overarching functions, with naming based on leading-edge protein functions and secondary considerations based on the constituent pathway names. The main groups containing ten or more pathways were labeled: Proteasome, MAPK and PI3K/AKT, Complement System, Lysosomal Glycan Metabolism, Rho GTPase, Ribosome, and Other.

### Network Development

A network visualization was generated by organizing the GSEA results using R and Cytoscape.^29^ Model nodes were represented as rectangles, and pathway nodes were represented as either upward-facing or downward-facing triangles, indicating positive or negative Normalized Enrichment Score (NES), respectively. Two types of edges were defined. The first edge type connected model nodes to all their significant pathways from gene set enrichment analysis. The second edge type connected pathways with a cosine similarity score greater than 0.2. Because greater emphasis was placed on leading-edge proteins than pathway names, each significant pathway for each model was retained as a unique node in the network. For example, the “Biocarta Proteasome Pathway” was significantly reduced for the duration of play model and significantly enriched for the RHI vs. low CTE, RHI vs. high CTE, and dementia models. However, all of these pathways were placed in different positions during cosine similarity score hierarchical clustering based on their unique leading-edge proteins. Therefore, they also occupied unique nodes in the final network, which was organized based on those cosine similarity scores. Within Cytoscape, the network was set to cluster based on the cosine similarity scores. Node colors were assigned based on hierarchical cluster group membership, and shapes were used to differentiate models, enriched pathways, and reduced pathways.

## Results

The number of brain donors included in each model and summary statistics of differential protein expression and gene set enrichment analysis are detailed in Table 2. Overall, most models included at least 80% of the brain donors, with the exception of the RHI vs. low CTE model.

### Early and Late CTE Changes Are Concordant

To characterize stage-specific proteomic changes, we first compared protein expression between CTE stages and RHI controls as the baseline. Analysis identified 58 significantly differentially expressed proteins (32 increased, 26 decreased; Fig. 2A) between the RHI and low CTE groups. Comparison of RHI and high CTE identified 262 significantly differentially expressed proteins (91 increased, 171 decreased; Fig. 2B).

**Figure 2.**
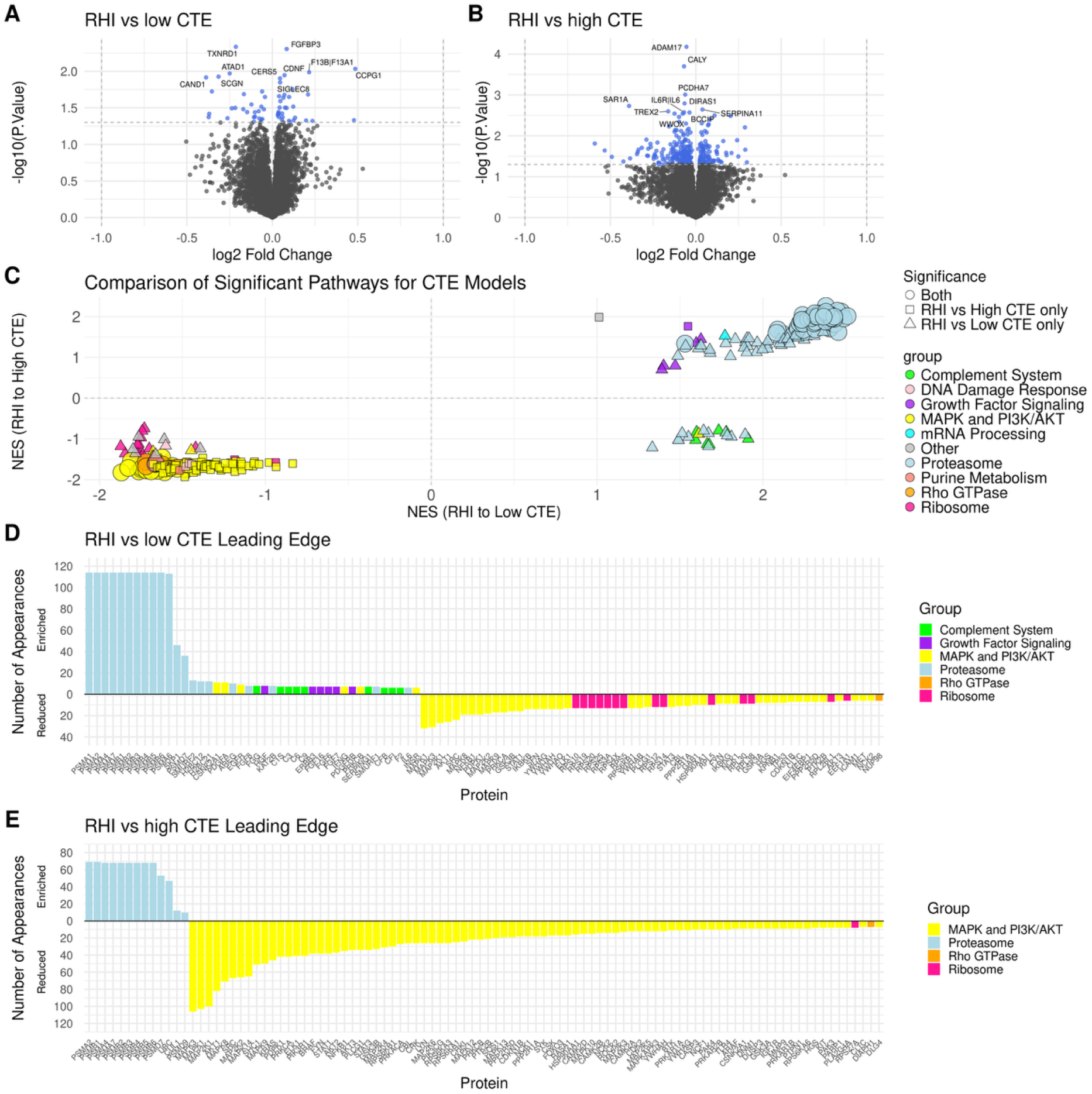
Proteome Changes Associated with Low CTE and High CTE Compared to RHI. (A) Volcano plot comparing RHI and low CTE groups. Significantly differentially expressed proteins with p<0.05 are shown in blue. (B) Volcano plot comparing RHI and high CTE groups. Significantly differentially expressed proteins with p<0.05 are shown in blue. (C) Scatter plot comparing NES values for all significant pathways in both CTE models, grouped based on hierarchical clustering. (D) Bar plot of the most frequently appearing leading-edge proteins for the RHI vs. low CTE model, showing the prevalence of proteasome subunit proteins (blue) in pathway analysis. (E) Bar plot of the most frequently appearing leading-edge proteins for the RHI vs. high CTE model, showing continued prevalence of proteasome subunit proteins (blue) as well as MAPK proteins (yellow). Bars above and below the zero line in D and E indicate the proteins were part of pathways that were enriched and reduced, respectively.

Pathway analysis revealed concordant trends between low CTE and high CTE relative to the RHI group. For the high CTE model, 92.4% of the significant pathways were either enriched proteasome pathways (31.4%) or reduced MAPK and PI3K/AKT signaling pathways (61.0%) (Fig. 2C). For the low CTE model, 79.2% of the significant pathways were either enriched proteasome pathways (59.9%) or reduced MAPK and PI3K/AKT signaling pathways (19.3%). An additional 6.7% of significant pathways in the low CTE model were reduced ribosome pathways (Fig. 2C).

Notably, enriched pathways related to the proteasome were identified in both models (Fig. 2C). These pathway alterations were primarily driven by changes to proteasome subunits identified in the leading-edge. Both low and high CTE models shared a common set of proteasome subunit proteins in the leading-edges of enriched proteasome pathways (Supplementary Fig. 3).

Proteasome subunits comprised the majority of leading-edge proteins in the significant pathways for both CTE models (Figs. 2D and 2E), based on the frequency of each protein’s appearance across all significant pathways. For the low CTE model, 12 of the 13 most frequently appearing proteins were proteasome subunits, with one additional proteasome-related protein. Proteasome subunit proteins also represented 10 of the 20 most frequently appearing proteins for the high CTE model.

The remaining 10 of the 20 most frequently appearing leading-edge proteins in the high CTE model were MAPK and PI3K/AKT proteins, highlighting the second largest grouping of pathways in the CTE models. The MAPK and PI3K/AKT signaling pathways are associated with cell signaling and affect processes such as cell growth, survival, and proliferation. These pathways had negative NES values in both CTE models (Fig. 2C). Further analysis of individual proteins of interest is detailed in Supplementary Fig. 2.

### Cortical Tau and Duration of play Have Distinct Features

The AT8 total and duration of play models identified 109 significant proteins (44 increased, 65 decreased; Fig. 3A) and 588 significant proteins (164 increased, 424 decreased; Fig. 3B), respectively. Consistent with the high CTE model, both models exhibited a greater proportion of downregulated than upregulated proteins. Specifically, 59.6% of significant proteins were downregulated in the AT8 total model, and 72.1% were downregulated in the duration of play model.

**Figure 3.**
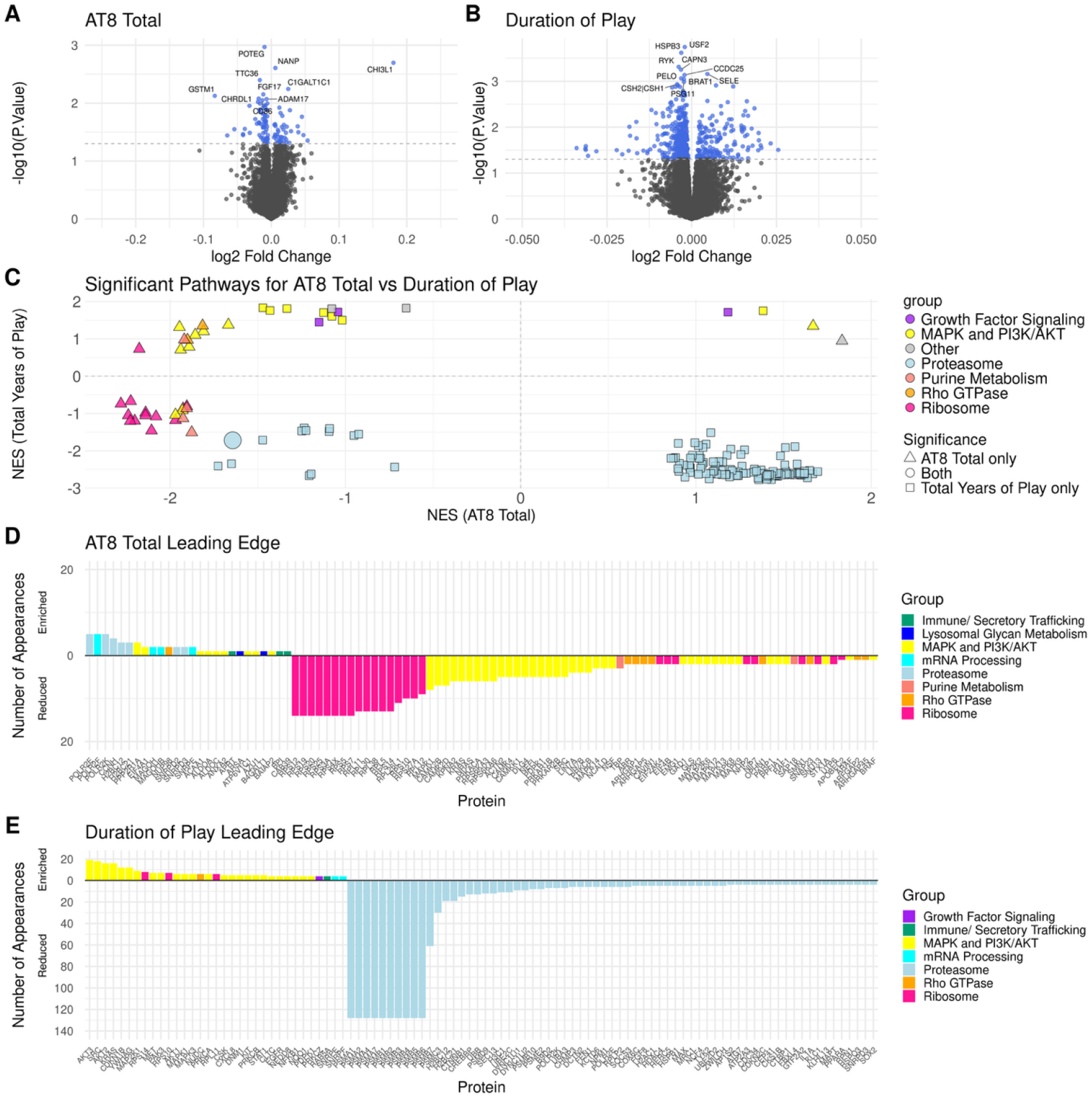
Proteome Changes Associated with Tau Burden and Duration of play. (A) Volcano plot for the AT8 total model. Significantly differentially expressed proteins with p<0.05 are shown in blue. (B) Volcano plot for the duration of play model. Significantly differentially expressed proteins with p<0.05 are shown in blue. (C) Scatter plot comparing NES values for all significant pathways in both models, with pathway groups named based on hierarchical clustering. (D) Bar plot of the most frequently appearing leading-edge proteins in the AT8 total model, displaying the reduction in ribosome proteins. (E) Bar plot of the most frequently appearing leading-edge proteins in the duration of play model showing reduction of proteasome subunit proteins (blue). Bars above and below the zero line in D and E indicate the proteins were part of pathways that were enriched and reduced, respectively.

Gene set enrichment analysis revealed notable differences between the AT8 total and duration of play models, suggesting distinct contributions to CTE progression. For the AT8 total model, tau burden was associated with a reduction in ribosomal proteins and pathways, particularly ribosome subunits. Ribosomal pathways constituted 14 of the 30 significant pathways for the AT8 model, forming the large pink cluster shown in Fig. 3C. One of these proteins, RPS12 (*P* = 0.028), was also significant in the AT8 total differential protein level analysis.

For the duration of play model, pathways with proteasome subunits in the leading-edge were reduced with increasing duration of play (Fig. 3C). Consistent with the CTE models, the duration of play model displayed altered proteasome pathways driven by proteasome subunits. Nine of these subunits were also significantly downregulated with increasing years of contact sports (Supplementary Fig. 3). Of the 141 significant pathways in this model, 129 were proteasome-grouped pathways containing these proteasome subunits. All of these pathways were reduced, indicating a strong correlation between duration of play and reduction in proteasome processes. Further analysis of individual proteins of interest is detailed in Supplementary Fig. 2.

In contrast to the CTE models, the most prevalent leading-edge proteins for the AT8 total model were ribosome subunit proteins (Fig. 3D). The 17 most frequently appearing proteins were all ribosome subunits, with the remaining three proteins in the top 20 being KRAS, MAPK1, and NRAS. KRAS and NRAS both clustered into the MAPK and PI3K/AKT Signaling Network group but were especially prevalent in synaptic-related pathways within this broader group. Further information regarding the ribosome subunit proteins is detailed in Supplementary Fig. 3.

The most prevalent leading-edge proteins for the duration of play model were proteasome subunit proteins (Fig. 3E). The 12 most frequently appearing proteins were all proteasome subunits. The next eight proteins were considerably less frequent and represented a mixture of proteasome-related proteins and MAPK and PI3K/AKT-related proteins. Further information regarding the proteasome subunit proteins is detailed in Supplementary Fig. 3.

### Cognitive Difficulty Scale and Dementia Associations are Concordant

Two Limma models evaluated protein changes associated with clinical measures of cognitive function and status: CDS and dementia status. Analysis identified 1,008 significant proteins for the dementia model (584 increased and 424 decreased; Fig. 4A) and 154 significant proteins for the CDS model (37 increased and 117 decreased; Fig. 4B).

**Figure 4.**
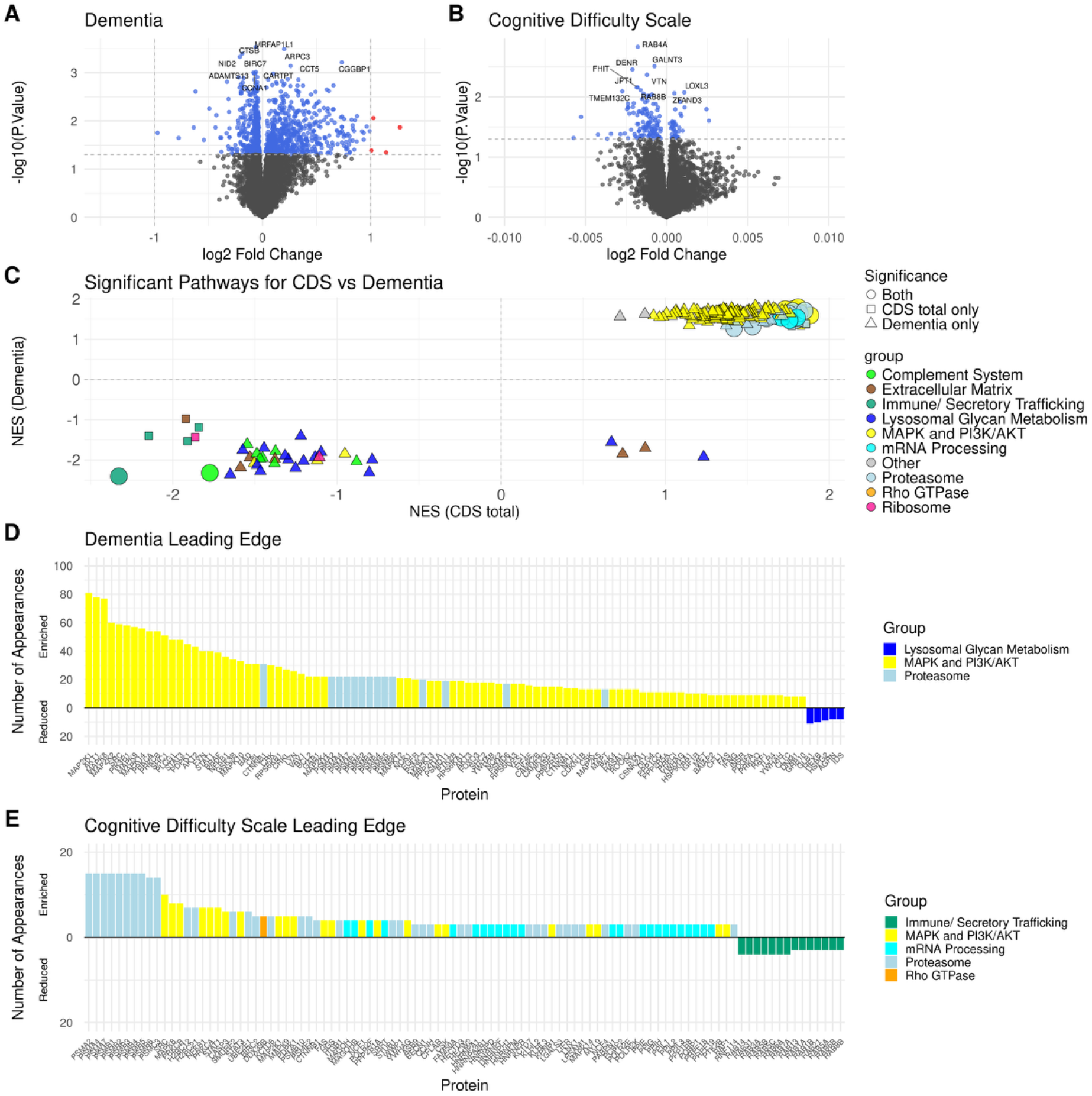
Proteome Changes Associated with Cognitive Dysfunction. (A) Volcano plot for the dementia model. Significantly differentially expressed proteins with p<0.05 are shown in blue; proteins with log2 fold change values greater than 1 are shown in red. (B) Volcano plot for the CDS model. Significantly differentially expressed proteins with p<0.05 are shown in blue. (C) Scatter plot comparing NES values for all significant pathways in both models, with pathway groups named based on hierarchical clustering. (D) Bar plot of the most frequently appearing leading-edge proteins in the dementia model. (E) Bar plot of the most frequently appearing leading-edge proteins in the CDS model. Bars above and below the zero line in D and E indicate the proteins were part of pathways that were enriched and reduced, respectively.

Pathway analysis revealed enrichment of proteasome pathways in both the CDS and dementia models (Fig. 4C), with 16 of 37 significant pathways for CDS and 20 of 215 significant pathways for dementia clustering to the proteasome group. These pathways contained the same leading-edge proteasome subunit proteins identified in the CTE models. The dementia model additionally contained enrichment in 148 of 215 pathways (68.8%) that clustered with the MAPK and PI3K/AKT Signaling Pathway group. The dementia model also demonstrated a reduction in the Lysosomal Glycan Metabolism group. Further analysis of significant pathways and groups is detailed in Supplementary Fig. 4, and individual proteins of interest are detailed in Supplementary Fig. 2

The leading-edge protein appearances within the dementia model were dominated by the MAPK and PI3K/AKT Signaling Pathway group (Fig. 4D). These proteins were related to cell signaling, proliferation, and survival. The leading-edge proteins for the CDS model were predominantly proteasome subunit proteins (Fig. 4E). The 10 most frequently appearing proteins for this model were all proteasome subunits, and an additional two proteasome subunits appeared among the top 20. The remaining proteins comprised a mixture of MAPK and PI3K/AKT Signaling Pathway group proteins and non-subunit proteins that were nevertheless associated with the proteasome group.

### Model Comparisons

Each of the models considered in this study provided insight into different aspects of CTE, from exposure risk factors to end-stage pathological endpoints. To elucidate how these results relate to each other, significant pathways from all six models were combined into a network diagram (Fig. 5). Briefly, significant pathways from each model were clustered based on the leading-edge genes driving the enrichment, where rectangular nodes represent models, upward- and downward-facing triangle nodes represent pathways, and edges represent significance or leading-edge overlap (see Methods). Groups labeled in Fig. 5 contain at least ten pathways. All other groups with five to ten pathways are detailed in Supplementary Fig. 4.

**Figure 5.**
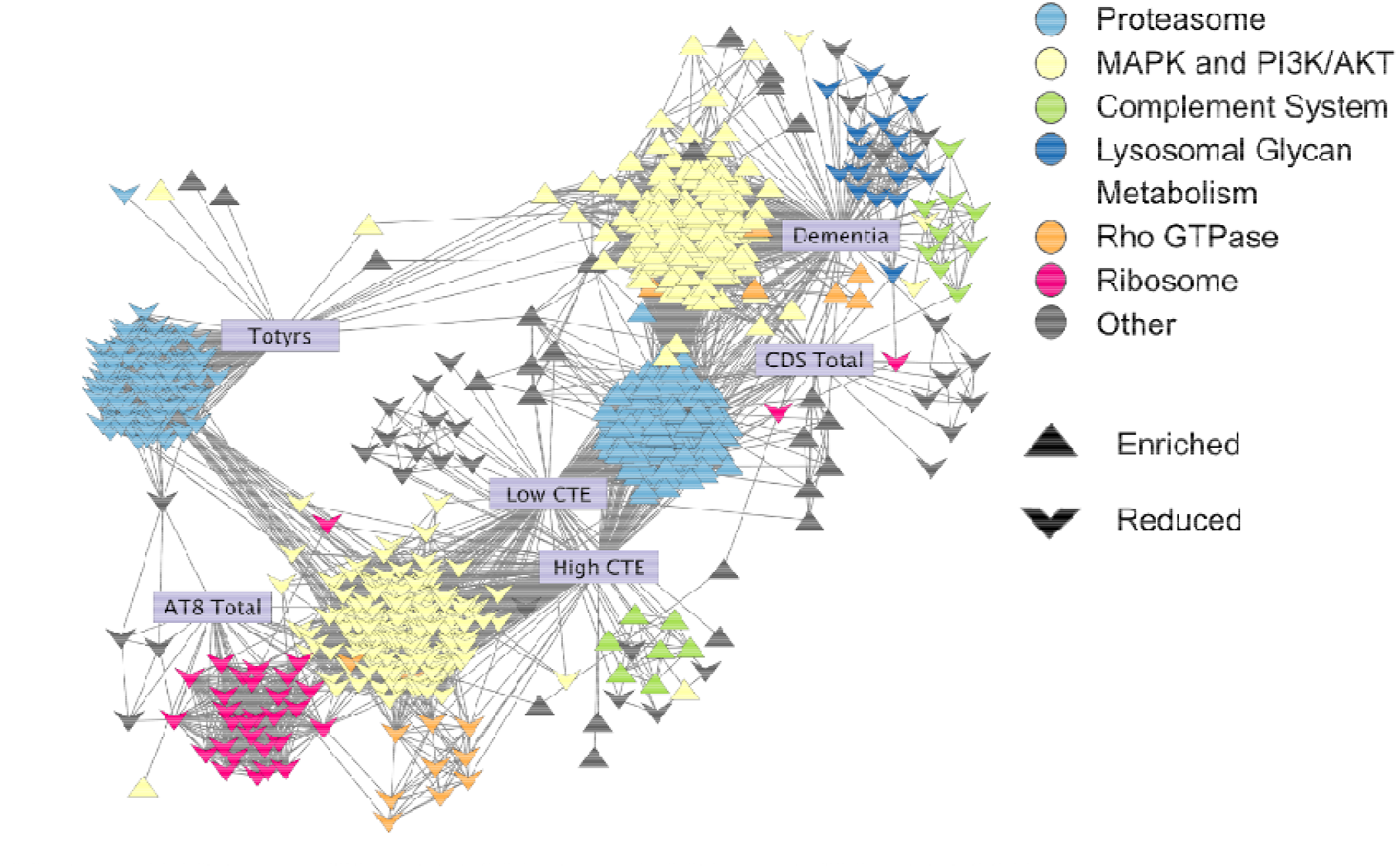
Synthesis of Models. Network of all significant pathways across all six models in this study. Purple rectangles denote the six models; upward-facing triangles denote enriched pathways; downward-facing triangles denote reduced pathways. Significant pathways are connected if their cosine similarity score is greater than 0.2, and pathways are also clustered based on this value. Color coding denotes cluster membership, which is based on cosine similarity score hierarchical clustering.

Overall, the most pronounced effects were observed in two main pathway groups: the proteasome pathways and the MAPK and PI3K/AKT signaling pathways. For the proteasome group, the CTE and cognitive test models demonstrated enrichment in proteasome pathways, while the duration of play model demonstrated a strong reduction. Proteasome-associated pathways accounted for 346 of 825 (41.9%) total significant pathways across all six models, with 129 of those 346 pathways representing reductions in the duration of play model.

The alterations in MAPK and PI3K/AKT signaling pathways suggest broad changes to cell signaling processes with predicted effects on cell survival, growth, and proliferation. These pathways constituted 340 of 825 (41.2%) significant pathways across all models, with 176 reduced and 164 enriched. Of the enriched MAPK and PI3K/AKT signaling pathways, 90.2% originated from the dementia model, and 93.8% of the reduced pathways were from the CTE models. Collectively, 83% of significantly altered pathways across all six models fell in either the proteasome or MAPK and PI3K/AKT groups.

The remaining 17% of significant pathways represented several smaller groups, with ribosome, complement system, Rho GTPase, and Lysosomal Glycan Metabolism constituting the majority. The ribosome group accounted for 31 of the 825 significant pathways. The complement system and Rho GTPase groups comprised 16 and 17 of the 825 significant pathways, respectively, with a mix of enriched and reduced pathways in each. Finally, the reduction observed in the Lysosomal Glycan Metabolism group was exclusive to the dementia model and accounted for 16 of the 825 pathways.

## Discussion

This study characterizes proteomic changes associated with CTE pathogenesis and key clinical and pathological features of the disease. The results suggest that several distinct biological pathways are altered in CTE, varying in association with RHI exposure, clinical endpoints, and pathological characteristics. Specifically, this study identifies close associations between the reduction of proteasome-related pathways, significant downregulation of proteasome subunit proteins, and increased years of contact sports play. An association between the reduction of ribosome-related pathways and tau burden was also identified through the AT8 total model. Finally, examining the CTE and dementia models revealed close associations with both proteasome and MAPK and PI3K/AKT pathways. Overall, the strongest associations with CTE and related models involved the proteasome and MAPK and PI3K/AKT pathways.

These results reveal an important role for the proteasome in CTE pathogenesis. Duration of play was associated with significant reductions in many of the proteasome’s alpha and beta subunits, denoted PSMA and PSMB. These subunits constitute the 20S catalytic core of the proteasome.^31^ Previous research has suggested that proteasome activity is linked to the expression of proteasome subunits.^32^ Therefore, in the context of these findings, a reduction in proteasome activity may be associated with duration of play.^32^ The observation that increased duration of play is associated with a reduction in proteasome-related pathways further supports this interpretation. However, the CTE models demonstrated enrichment of proteasome-related pathways without significant changes in individual proteasome subunits in the differential expression analysis. The initial enrichment of proteasome processes observed in the CTE models may indicate a compensatory response to the aggregation of pathological tau. However, the subsequent reduction of proteasome pathways in high CTE compared to low CTE may reflect eventual proteasome dysfunction, possibly when the system is overwhelmed by tau aggregation or its resulting oxidative stress. This transition from low to high CTE may correspond to the pattern observed in the duration of play model.

The proteasome dysregulation observed in CTE parallels findings in other neurodegenerative diseases, including Alzheimer’s disease (AD), Huntington’s disease, Parkinson’s disease, and amyotrophic lateral sclerosis (ALS).^31,33–37^ Previous work has demonstrated downregulation of specific proteasome subunits in each disease: PSMB5 in AD, PSMA7 in Parkinson’s disease, and PSMB3 in ALS.^29-33^ Moreover, proteasomal degradation can limit the propagation of pathological tau, a major driver of AD pathology.^34,35,37^ Taken together with these prior findings, proper proteasome formation and function may be critical in the body’s response to tauopathy and aberrant protein aggregation in general. This process appears to be enriched in low and high CTE, possibly as a compensatory mechanism.^34,35,37^ Our results also support the hypothesis that repeated head injuries associated with playing contact sports disrupt normal proteasome function in the brain, which could contribute to the aggregation of pathological tau and progression of CTE.

Consistent with the CTE models, the leading-edge proteins in the dementia model’s significant pathways were proteasome subunits, although these proteins were not significant in the differential expression analysis. However, several ubiquitin-associated proteins, including UBE2F, UBE2H, and UBE2S, were significantly upregulated in the dementia model. A recent study in AD suggested that when proteasome function is compromised, a compensatory increase in ubiquitin-associated protein levels occurs as a result.^38^ This may explain the observed increase in ubiquitin-associated proteins coinciding with dementia status, which is a symptom of CTE that predominantly manifests in later stages when the proteasome appears to be more dysregulated based on our results.

Other significantly altered pathways were more challenging to interpret than the proteasome group. Pathways in this group relate to diverse functions, including synaptic function, MAPK, PI3K, mTOR and growth factor signaling, immune/inflammatory signaling, hormone signaling, stress response, cell cycle and proliferation, metabolic regulation, and oncogenic signaling.^31^ These pathways were reduced primarily in the CTE models and enriched primarily in the dementia model. Both CTE models contained significant pathways related to MAPK and PI3K processes. However, the low CTE pathways included processes related to adaptive stress responses (8 of 37 pathways), while the high CTE pathways included processes related to immune activation (27 of 128 pathways) and synaptic and vascular dysfunction (12 of 128 pathways), suggesting a transition from stress response to dysfunction. Among the specific cell signaling proteins significantly reduced in the context of CTE, CDK5 was reduced in the comparison of RHI vs. high CTE, and MAPK3 (ERK1) was reduced in both the RHI vs. low CTE and RHI vs. high CTE models. Previous research has demonstrated that changes to CDK5 correlate with impaired axonal guidance, suggesting these cell signaling changes may contribute to the synaptic deficits observed in CTE pathology.^9^ The pathways enriched in the dementia model indicate a shift from normal neuronal signaling toward an injury-response state characterized by chronic neuroinflammation, aberrant receptor tyrosine kinase signaling, altered adhesion and migration, oxidative stress, and dysregulated plasticity.

Tau burden in the cortical gyrus was found to be closely associated with a reduction of ribosome-associated protein pathways. This finding, together with the significant reduction in RPS12 (a ribosome subunit) with increasing tau burden, supports recent literature highlighting important connections between ptau and ribosomal activity in AD.^32–34^ More specifically, human studies examining AD demonstrated that pathological tau associated closely with the ribosome and reduced translation levels, and that nucleolar chaperone proteins and RNA Polymerase I were reduced in AD pathology.^32,33^ A mouse study examining AD models with hTau mutations further demonstrated that pathological tau reduced ribosome biogenesis and protein synthesis.^39^ These studies suggest that the ability of tau aggregates to compromise protein synthesis likely results in disruptions to synaptic plasticity and general proteostasis, ultimately contributing to brain atrophy.^39,40^ A significant increase in CHI3L1 levels with increased tau burden was also observed, which is notable given that CHI3L1 is considered a key mediator of inflammatory processes.^41^

Consistent with previous studies of CTE, we observed signs of neuroimmune and glial changes associated with CTE pathology, both in terms of altered cytokine levels and changes in the levels of GFAP, a marker of reactive astrocytes.^42^ In the comparison of RHI and high CTE, CCL22, CXCL13, and IL6 were significantly reduced, while GFAP levels were increased. It is notable that increased GFAP levels were also associated with tau burden and duration of play.^43^

Several notable changes in integrin proteins and growth factors were also observed. Significant changes in ITGB1 and ITGB5 were identified when comparing RHI and low CTE, and significant changes in ITGB3 and ITGB6 were identified when comparing RHI and high CTE. Changes in integrin expression have also been noted in recent single-nucleus RNA sequencing experiments analyzing transcriptional changes associated with CTE.^3^ Previous studies have established that integrins play an important role in maintaining the integrity of the blood–brain barrier. Therefore, the observed disruptions of integrin expression in CTE may help to explain the heightened infiltration of T cells into the brain noted in prior studies.^44,45^ This would be consistent with our observation that CD247, a T-cell marker, was significantly elevated in high CTE compared to RHI cases.^46^ Regarding significant changes in growth factor levels, CDS scores were associated with a decrease in BDNF, which is consistent with the established roles of BDNF in neuronal survival and synaptic health.^47,48^

It is important to acknowledge several limitations of this study. First, no female cases were included due to a lack of available samples. This reflects the fact that study samples were dominated by American football players, and females have historically not had opportunities to participate in this sport. Second, the available brains were disproportionately in the later stages of CTE: 58 brain donors in the RHI group, 33 in the low CTE group, and 113 in the high CTE group. This distribution reflects the ascertainment bias inherent in brain donation, as individuals are more likely to donate if they exhibited more severe symptoms. Third, the AT8 total feature evaluates abnormal tau in only one specific brain region. While the dorsolateral frontal cortex is an important area, as it is one of the first locations where CTE pathology is identified, it cannot capture the full extent of abnormal tau throughout the brain.^49^

Taken together, these findings advance our understanding of CTE and suggest important correlations between duration of play, proteasome dysfunction, and the progression to later stages of CTE, as well as the effects of pathological tau on the ribosome. This study also identifies potential shared mechanisms between CTE and other neurodegenerative diseases, including Alzheimer’s disease.^33–35^ By providing further characterization of the specific protein changes that occur across different stages of CTE, this study may help identify important clinical targets and disease protein biomarkers in the brain, a major goal of CTE research.^50^ These results also help identify which specific proteins and pathways are most strongly correlated with factors such as duration of play and functional outcomes including cognitive decline and dementia.

## Data Availability

All data and corresponding code used in this study can be found in our OSF project and GitHub.

OSF Link: https://osf.io/5xach

GitHub Link: https://github.com/BU-Neuromics/cte_proteomics_2026

## Acknowledgements

We extend thanks to Simon Dillon for producing our SomaScan 7k data.

## Supplementary Material

Supplementary material is available online at https://osf.io/5xach.

