## Supplemental Figures and Tables for "Proteomic Analysis of Human Chronic Traumatic Encephalopathy Brain Implicates Proteasome and Ribosome Dysfunction in Disease Progression"

Changes to the Proteasome System in CTE Implicated by Proteome Analysis – Figures

**Figure 1. SomaScan 7k Proteomics Filtering and Preprocessing**

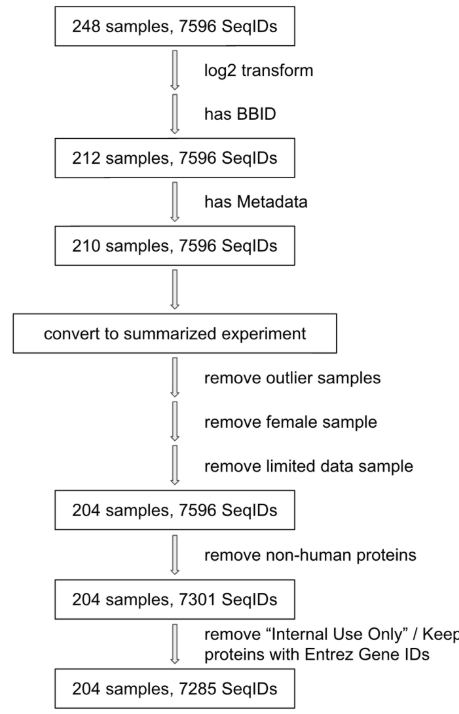

Flow Chart of SomaScan 7k Proteomics Filtering. Boxes represent updated numbers of samples and proteins included in the study. Arrows detail the filtering and transformation steps taken to get to our final data set used in the study.

**Figure 2. Proteome Changes Associated with Low CTE and High CTE Compared to RHI**

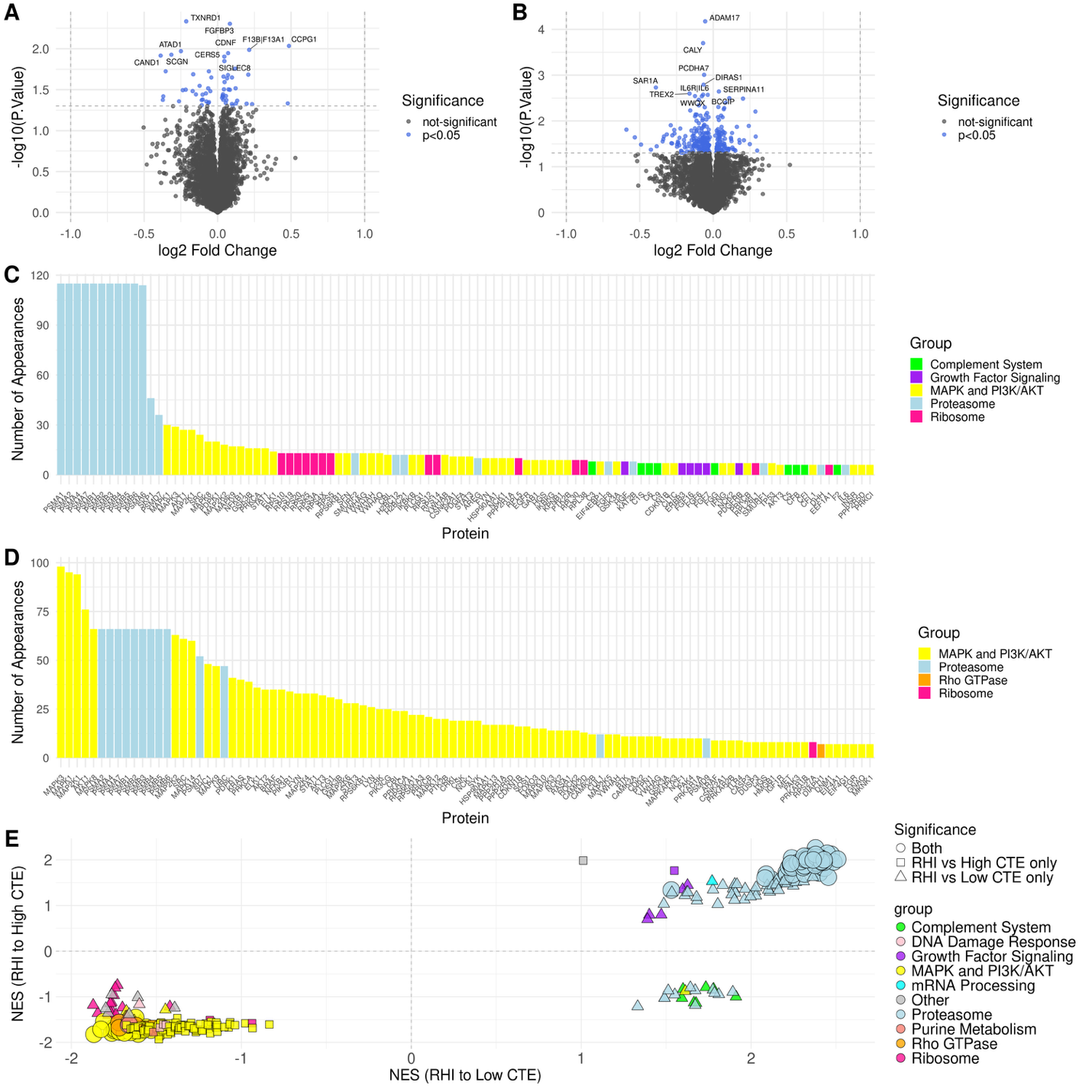

a. Volcano plot for RHI vs low CTE detailing significantly differentially expressed proteins in blue, b. Volcano plot for RHI vs high CTE detailing significantly expressed proteins in blue, c. Bar plot of most common leading-edge proteins for the RHI vs low CTE model showcase prevalence of proteasome subunit proteins (blue) in pathway analysis, d. Bar plot of most common leading-edge proteins for the RHI vs high CTE model showcase continued prevalence of proteasome subunit proteins (blue) in pathway analysis as well as prevalence of MAPK proteins (yellow), e. Scatterplot comparing NES values for all significant pathways for both RHI vs low CTE and RHI vs high CTE grouped based on hierarchical clustering.

**Figure 3: Proteome Changes Associated with Tau Burden and Total Years of Violent Sports Play**

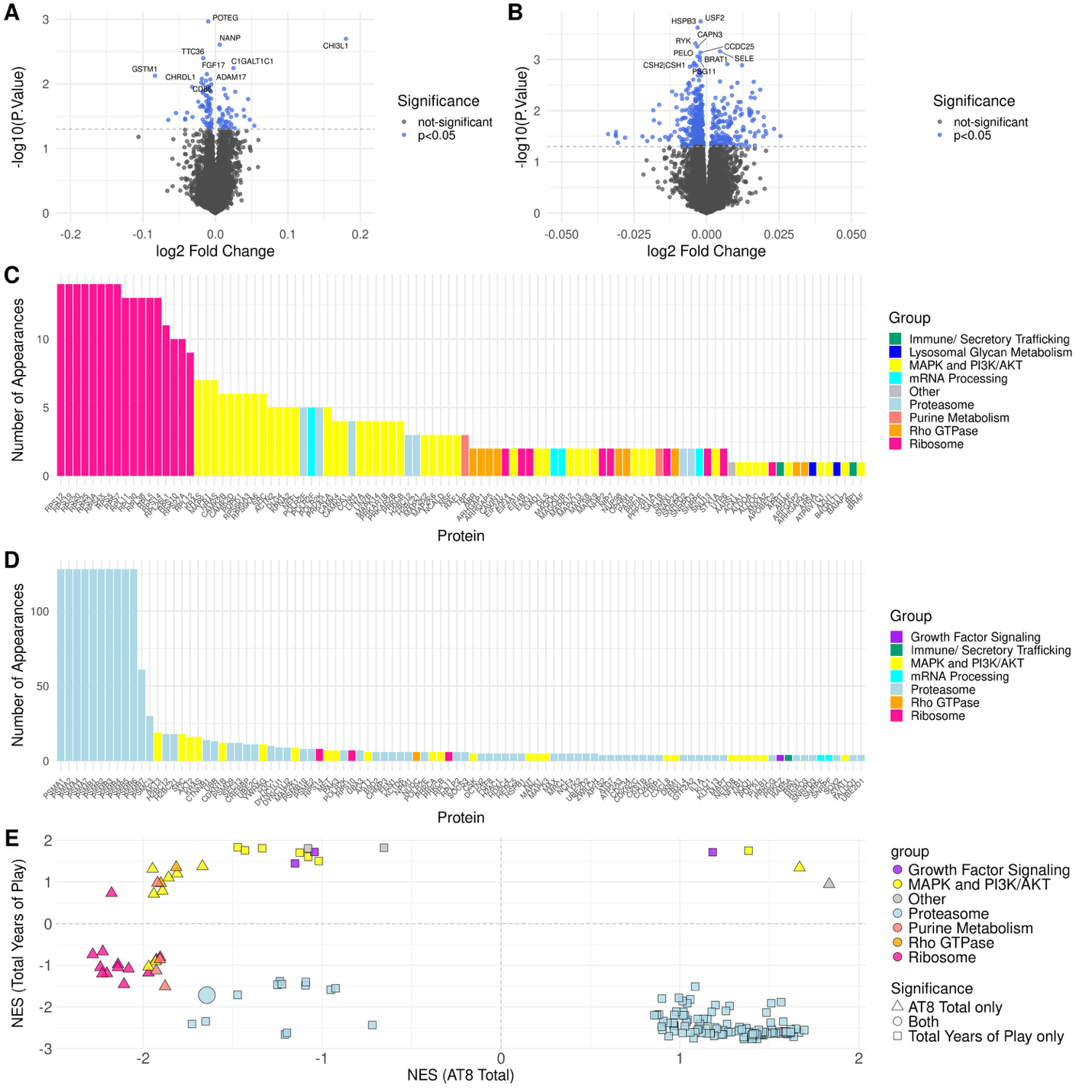

a.  Volcano plot for the AT8 total model detailing significantly differentially expressed proteins in blue, b. Volcano plot for the total years of contact sports play model detailing significantly differentially expressed proteins in blue, c. Bar plot of most common proteins in the leading-edge of significant pathways for the AT8 total model, d. Bar plot of most common proteins in the leading-edge of significant pathways for the total years of contact sports play model, e. Scatterplot comparing NES values for all significant pathways in the total years of contact sports play model and the AT8 total model with pathway groups named based on hierarchical clustering.

**Figure 4: Proteome Changes Associated with Cognitive or Functional Decline**

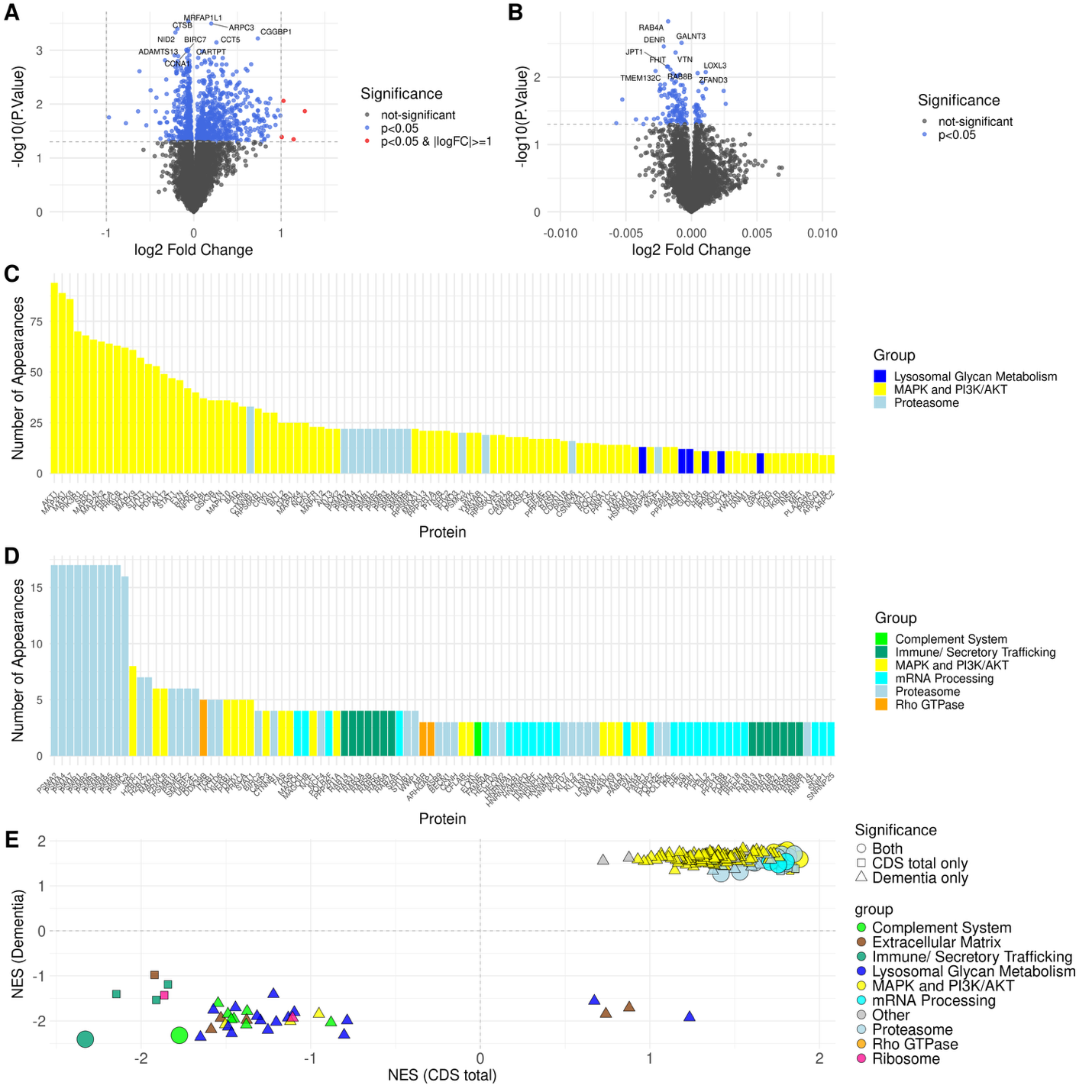

a. Volcano plot for the Dementia model detailing significantly differentially expressed proteins in blue and differentially expressed proteins with log2 fold change values greater than 1 in red, b. Volcano plot for the CDS total model detailing significantly differentially expressed proteins in blue, c. Bar plot of most common proteins in the leading-edge of significant pathways in the Dementia model. d. Bar plot of most common proteins in the leading-edge of significant pathways in the CDS total model. e. Scatterplot comparing NES values for all significant pathways in the CDS total model and the Dementia model with pathway groups named based on hierarchical clustering. 
**Figure 5. Synthesis of Models**

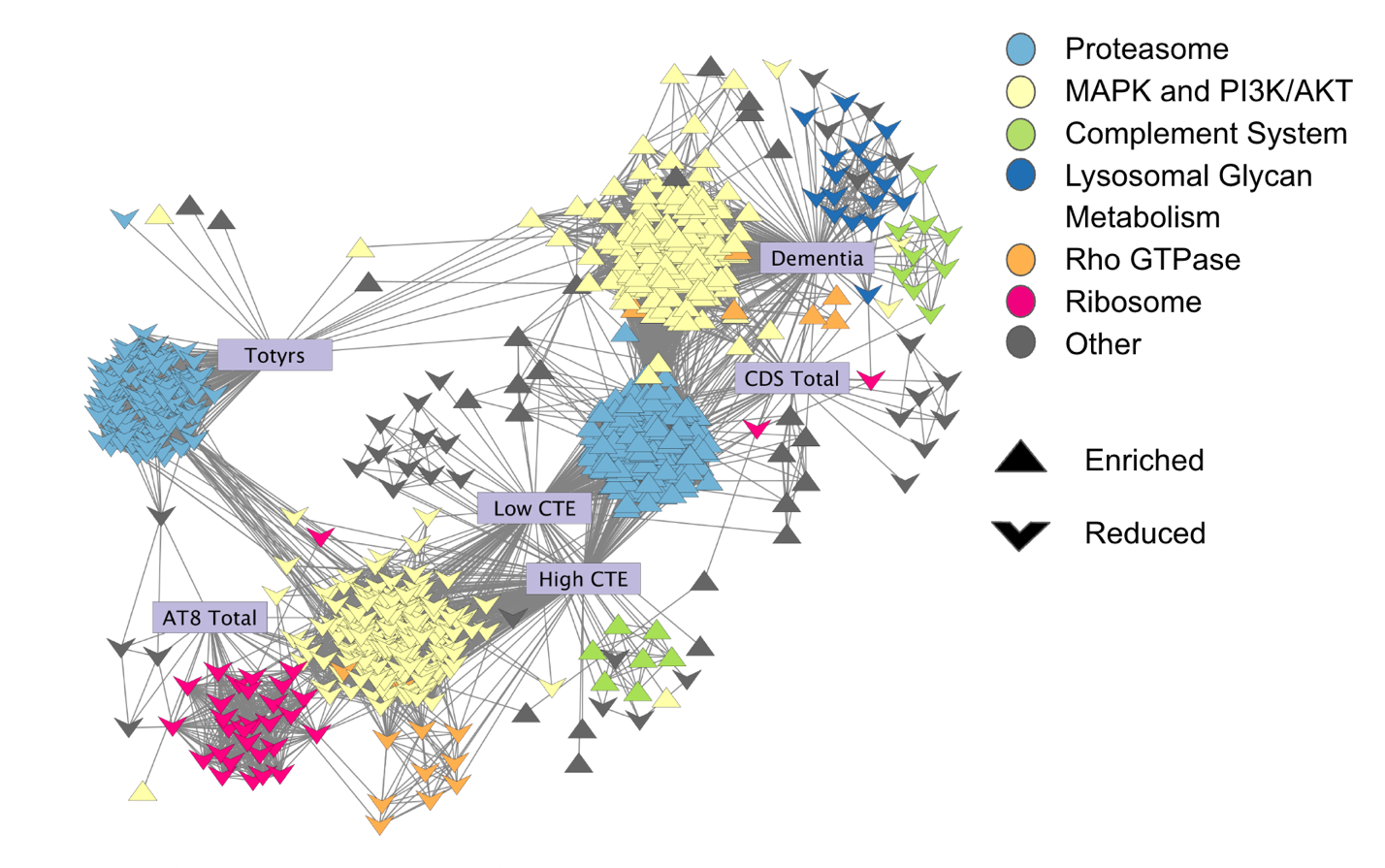
Network of all significant pathways across all six models in this study. Purple rectangles denote the six models; upwards facing triangles denote the enriched pathways, and downwards facing triangles denote reduced pathways. Significant pathways are connected if their cosine similarity score is greater than 0.2, and pathways are also clustered based on this value. Color coding of the pathways denotes which cluster of pathways they belong to, which is also based on the cosine similarity score hierarchical clustering.

**Additional or Supplemental Figures**

**Supplemental Figure 1.**

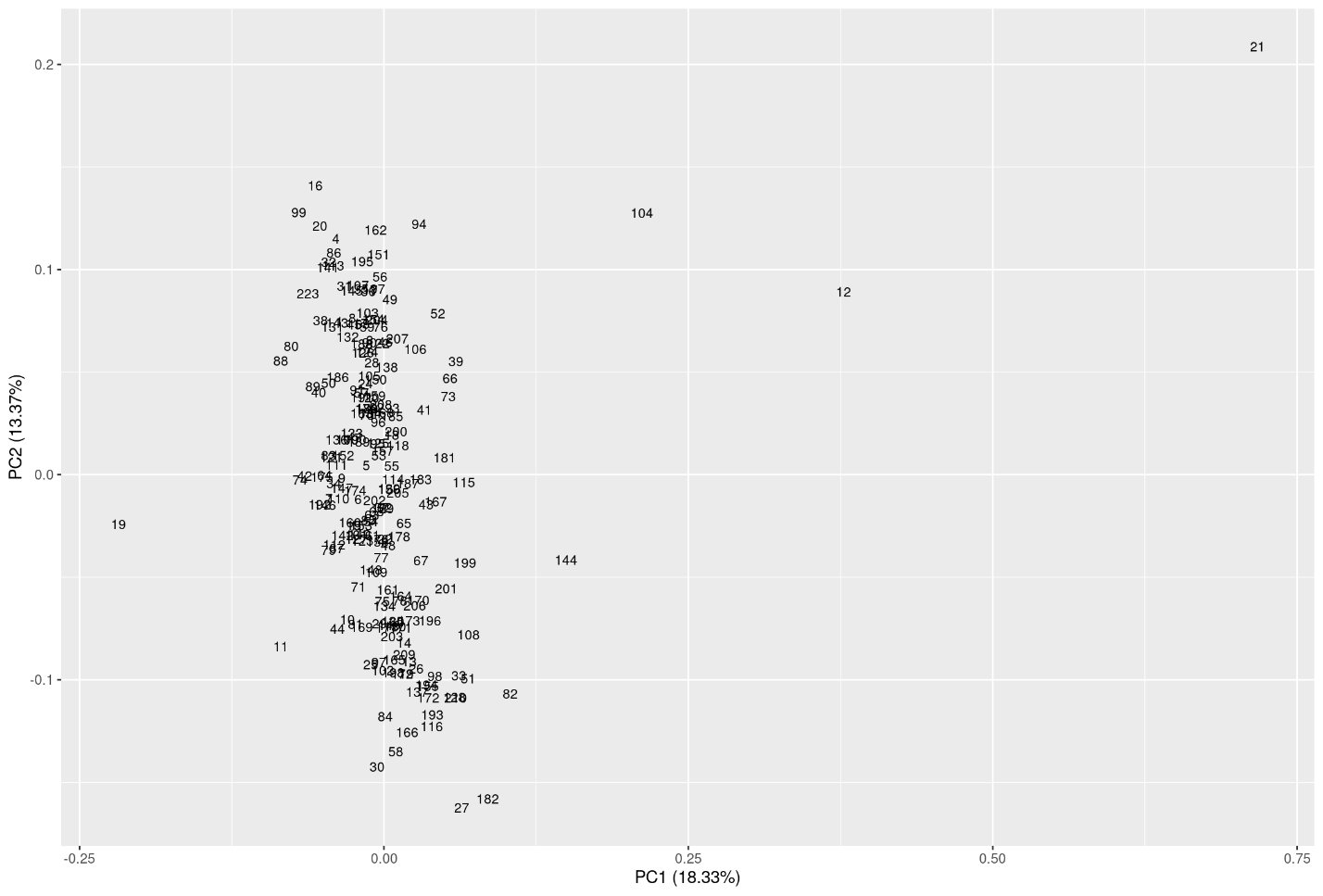

**Supplemental Figure 1. PCA plot of all SomaScan protein samples identifies three outlier samples.** Samples 12, 19, 21, and 104 were identified as outliers with standard deviations greater than 3 and removed from the data set using the PCA plot displayed above.

**Supplemental Figure 2:**

 Notable Significant Proteins by Model

Proteins of interest by model are listed here along with their log fold change (LFC) and p-values. In models examining CTE, a positive value indicates a significant increase associated with CTE. In the case of factors such as tau burden or dementia, a positive value indicates an increase in protein levels with the incidence of these factors.

| RHI vs High CTE | | | |
| --- | --- | --- | --- |
| **Proteasome-Associated Proteins** | | | |
| UBA3 | LFC = -0.089   p = 0.019 |  |  |
| **Integrin Proteins** | | | |
| ITGB3 | LFC = -0.125   p = 0.014 | ITGB6 | LFC = 0.033   p = 0.015 |
| **Inflammation-Associated Proteins** | | | |
| GFAP | LFC = 0.110   p = 0.003 | CCL3L1 | LFC = -0.051   p = 0.018 |
| CCL22 | LFC = -0.027   p = 0.041 | TLR2 | LFC = -0.037   p = 0.040 |
| IL6 | LFC = -0.069   p = 0.002 | CXCL13 | LFC = -0.082   p = 0.032 |
| CD247 | LFC = 0.122  p = 0.025 |  |  |
| **Cell-Signaling and MAPK Proteins** | | | |
| CDK5 | LFC = -0.178   p = 0.047 | MAPK3 | LFC = -0.087  p = 0.028 |
| **Synaptic Proteins** | | | |
| SLITRK6 | LFC = -0.031   p = 0.033 | LRRC20 | LFC = 0.079  p = 0.044 |
| LRRC3B | LFC = 0.072  p = 0.006 | LRRTM4 | LFC = 0.035  p = 0.005 |
| RHI vs Low CTE | | | |
| **Proteasome-Associated Proteins** | | | |
| USP28 | LFC = 0.064   p = 0.023 |  |  |
| **Integrin Proteins** | | | |
| ITGB5 | LFC = 0.037   p = 0.038 | ITGB1 | LFC = -0.080   p = 0.035 |
| **Inflammation-Associated Proteins** | | | |
| CXCL2 | LFC = 0.083   p = 0.032 | IFNB1 | LFC = 0.031   p = 0.040 |
| **Cell-Signaling and MAPK Proteins** | | | |
| MAPK3 | LFC = -0.108   p = 0.049 |  |  |
| **Synaptic Proteins** | | | |
| LRRC3B | LFC = 0.074  p = 0.031 |  |  |

| Total Years of Contact Sports | | | |
| --- | --- | --- | --- |
| **Proteasome-Associated Proteins** | | | |
| PSMA2 | LFC = -0.031   p = 0.029 | PSMA7 | LFC = -0.031   p = 0.026 |
| PSMB1 | LFC = -0.034   p = 0.029 | PSMB3 | LFC = -0.016   p = 0.018 |
| PSMB4 | LFC = -0.028   p = 0.034 | PSMB5 | LFC = -0.031   p = 0.031 |
| PSMB7 | LFC = -0.005   p = 0.018 | RPN1 | LFC = -0.009   p = 0.035 |
| UBE2D1 | LFC = -0.003   p = 0.004 | UBE2J1 | LFC = -0.003   p = 0.005 |
| UBE2D2 | LFC = -0.004   p = 0.006 | KEAP1 | LFC = -0.003   p = 0.018 |
| **Inflammation-Associated Proteins** | | | |
| GFAP | LFC = 0.006   p = 0.009 | IL6 | LFC = 0.004   p = 0.031 |
| IL1RL1 | LFC = 0.023   p = 0.023 | CCL28 | LFC = 0.007   p = 0.035 |
| CXCL2 | LFC = 0.002   p = 0.014 | CX3CL1 | LFC = -0.010   p = 0.015 |
| IFNL4 | LFC = -0.002   p = 0.046 | IL13 | LFC = -0.002   p = 0.003 |
| IFNA5 | LFC = -0.002   p = 0.007 | IL17C | LFC = -0.002   p = 0.009 |
| CXCL8 | LFC = 0.011  p = 0.003 |  |  |
| **Synaptic Proteins** | | | |
| SLITRK1 | LFC = -0.008   p = 0.014 | SLITRK6 | LFC = 0.002   p = 0.048 |
| SNAP25 | LFC = -0.002   p = 0.007 |  |  |
| **Integrin Proteins** | | | |
| ITGB5 | LFC = -0.007   p = 0.029 |  |  |

| Tau Burden | | | |
| --- | --- | --- | --- |
| **Proteasome-Associated Proteins** | | | |
| UBE2D3 | LFC = 0.054   p = 0.044 |  |  |
| **Ribosome-Associated Proteins** | | | |
| RPS12 | LFC = -0.039   p = 0.028 |  |  |
| **Inflammation-Associated Proteins** | | | |
| GFAP | LFC = 0.014   p = 0.049 | CHI3L1 | LFC = 0.180   p = 0.002 |
| IFNA14 | LFC = 0.006   p = 0.025 | CCL16 | LFC = -0.012   p = 0.019 |
| IFNA10 | LFC = 0.006   p = 0.048 |  |  |
| ￼**Synaptic Proteins** | ￼￼ | ￼￼ | ￼￼ |
| ￼SNAP23 | ￼LFC = -0.011  p = 0.033 | ￼LRRC3B | ￼LFC = 0.010  p = 0.038 |

| Dementia | | | |
| --- | --- | --- | --- |
| **Proteasome-Associated Proteins** | | | |
| UBA2 | LFC = 0.569   p = 0.005 | UBA6 | LFC = 0.800   p = 0.031 |
| UBA7 | LFC = 0.176   p = 0.025 | UBAC1 | LFC = 0.261   p = 0.019 |
| UBE2F | LFC = 0.213   p = 0.017 | UBE2H | LFC = 0.213   p = 0.039 |
| UBE2S | LFC = 0.053   p = 0.026 | UBE2T | LFC = 0.143   p = 0.048 |
| UBE2Q2 | LFC = 0.098   p = 0.013 | UBE4A | LFC = 0.412   p = 0.037 |
| USP4 | LFC = 0.509   p = 0.030 | USP11 | LFC = -0.177   p = 0.036 |
| USP22 | LFC = 0.290   p = 0.047 | USP25 | LFC = 0.241   p = 0.015 |
| UBL4A | LFC = -0.305   p = 0.022 | UBE2B | LFC = -0.045   p = 0.036 |
| UBE2L3 | LFC = -0.046   p = 0.048 | PSMD4 | LFC = 0.240   p = 0.008 |
| PSME3 | LFC = -0.041   p = 0.038 | UBB | LFC = -0.435   p = 0.014 |
| UBC | LFC = -0.235   p = 0.014 | CUL9 | LFC = -0.052   p = 0.016 |
| UBE2Z | LFC = -0.176   p = 0.007 |  |  |
| **Inflammation-Associated Proteins** | | | |
| CXCL2 | LFC = -0.043   p = 0.046 | CXCL9 | LFC = -0.117   p = 0.004 |
| CCL7 | LFC = -0.055   p = 0.013 | CCL22 | LFC = 0.035   p = 0.037 |
| CCL23 | LFC = -0.054   p = 0.029 | CCL8 | LFC = -0.070   p = 0.022 |
| C1QC | LFC = -0.043   p = 0.046 | IL5 | LFC = -0.054   p = 0.007 |
| IL12B | LFC = -0.032   p = 0.009 | IL1B | LFC = 0.101   p = 0.013 |
| IFNA10 | LFC = -0.045   p = 0.009 | IL17A | LFC = -0.039   p = 0.032 |
| **Synaptic Proteins** | | | |
| ROBO3 | LFC = -0.045   p = 0.035 | SYT12 | LFC = 0.430   p = 0.007 |
| **Integrin Proteins** | | | |
| ITGB1 | LFC = -0.066   p = 0.018 | ITGB2 | LFC = -0.132   p = 0.034 |
| **Cell-Signaling and MAPK Proteins** | | | |
| MAPK8 | LFC = 0.646   p = 0.028 | MAPK9 | LFC = 0.723   p = 0.017 |
| MAPK10 | LFC = 0.147   p = 0.007 | MAPK14 | LFC = 0.253   p = 0.040 |
| PRKCD | LFC = -0.040   p = 0.020 |  |  |

| FAQ score | | | |
| --- | --- | --- | --- |
| **Proteasome-Associated Proteins** | | | |
| PSMB9 | LFC = -0.004   p = 0.014 | PSMG2 | LFC = -0.004   p = 0.012 |
| UBE2L6 | LFC = -0.003   p = 0.022 | UBE2D2 | LFC = -0.003   p = 0.029 |
| UBE2Z | LFC = -0.006   p = 0.029 | KEAP1 | LFC = -0.002   p = 0.023 |
| PSME2 | LFC = -0.011   p = 0.026 |  |  |
| **Growth Factor Proteins** | | | |
| IGF1 | LFC = -0.003   p = 0.044 |  |  |
| CDS score | | | |
| **Proteasome-Associated Proteins** | | | |
| PSME3 | LFC = -0.001   p = 0.015 | UBE2V2 | LFC = -0.001   p = 0.027 |
| UBE2K | LFC = -0.001   p = 0.034 | UBE2A | LFC = 0.0004   p = 0.036 |
| **Growth Factor Proteins** | | | |
| BDNF | LFC = -0.001   p = 0.017 |  |  |

**Supplementary Figure 3:**

**Leading-Edge Proteasome Proteins**

| Proteasome Proteins | Model | Pvalue | LogFC |
| --- | --- | --- | --- |
| PSMB4 | RHI vs CTE Low | 0.379 | 0.253 |
|  | RHI vs CTE High | 0.504 | 0.143 |
|  | Dementia | 0.057 | 0.517 |
|  | CDS Total | 0.283 | 0.003 |
|  | Total Years of Play | 0.034* | -0.028 |
| PSMB2 | RHI vs CTE Low | 0.253 | 0.390 |
|  | RHI vs CTE High | 0.193 | 0.310 |
|  | Dementia | 0.137 | 0.455 |
|  | CDS Total | 0.534 | 0.002 |
|  | Total Years of Play | 0.042* | -0.031 |
| PSMA2 | RHI vs CTE Low | 0.221 | 0.398 |
|  | RHI vs CTE High | 0.300 | 0.238 |
|  | Dementia | 0.213 | 0.368 |
|  | CDS Total | 0.543 | 0.002 |
|  | Total Years of Play | 0.029* | -0.031 |
| PSMB1 | RHI vs CTE Low | 0.366 | 0.300 |
|  | RHI vs CTE High | 0.385 | 0.215 |
|  | Dementia | 0.151 | 0.455 |
|  | CDS Total | 0.593 | 0.002 |
|  | Total Years of Play | 0.029* | -0.034 |
| PSMB3 | RHI vs CTE Low | 0.270 | 0.173 |
|  | RHI vs CTE High | 0.484 | 0.076 |
|  | Dementia | 0.361 | 0.129 |
|  | CDS Total | 0.389 | 0.001 |
|  | Total Years of Play | 0.018* | -0.016 |
| PSMB6 | RHI vs CTE Low | 0.180 | 0.191 |
|  | RHI vs CTE High | 0.414 | 0.087 |
|  | Dementia | 0.366 | 0.123 |
|  | CDS Total | 0.580 | 0.001 |
|  | Total Years of Play | 0.052 | -0.013 |
| PSMA7 | RHI vs CTE Low | 0.287 | 0.335 |
|  | RHI vs CTE High | 0.357 | 0.204 |
|  | Dementia | 0.169 | 0.394 |
|  | CDS Total | 0.459 | 0.002 |
|  | Total Years of Play | 0.026* | -0.031 |
| PSMA4 | RHI vs CTE Low | 0.264 | 0.259 |
|  | RHI vs CTE High | 0.450 | 0.113 |
|  | Dementia | 0.274 | 0.231 |
|  | CDS Total | 0.396 | 0.002 |
|  | Total Years of Play | 0.031* | -0.022 |
| PSMB5 | RHI vs CTE Low | 0.330 | 0.317 |
|  | RHI vs CTE High | 0.288 | 0.245 |
|  | Dementia | 0.150 | 0.425 |
|  | CDS Total | 0.472 | 0.002 |
|  | Total Years of Play | 0.031* | -0.031 |
| PSMC3 | Dementia | 0.189 | 0.127 |
|  | CDS Total | 0.143 | 0.001 |
|  | Total Year of Play | 0.409 | -0.004 |
| PSMB7 | Total Years of Play | 0.018* | -0.005 |
| PSMA1 | Total Years of Play | 0.074 | -0.008 |

**Leading-Edge Ribosome Proteins**

| Ribosome Proteins | Model | Pvalue | LogFC |
| --- | --- | --- | --- |
| RPL5 | AT8 Total | 0.406 | -0.032 |
| RPL26L1 | AT8 Total | 0.557 | -0.014 |
| RPL38 | AT8 Total | 0.448 | -0.009 |
| RPL30 | AT8 Total | 0.468 | -0.023 |
| RPL11 | AT8 Total | 0.198 | -0.012 |
| RPL12 | AT8 Total | 0.694 | -0.006 |
| RPS14 | AT8 Total | 0.183 | -0.026 |
| RPS10 | AT8 Total | 0.631 | -0.006 |
| RPS12 | AT8 Total | 0.028* | -0.039 |
| RPS7 | AT8 Total | 0.198 | -0.020 |
| RPS20 | AT8 Total | 0.191 | -0.036 |
| RPS4X | AT8 Total | 0.272 | -0.025 |
| RPS19 | AT8 Total | 0.463 | -0.025 |
| RPS5 | AT8 Total | 0.536 | -0.009 |
| RPS25 | AT8 Total | 0.180 | -0.013 |
| RPS27A | AT8 Total | 0.553 | -0.006 |
| RPS3A | AT8 Total | 0.409 | -0.011 |

**Supplemental Figure 4.**

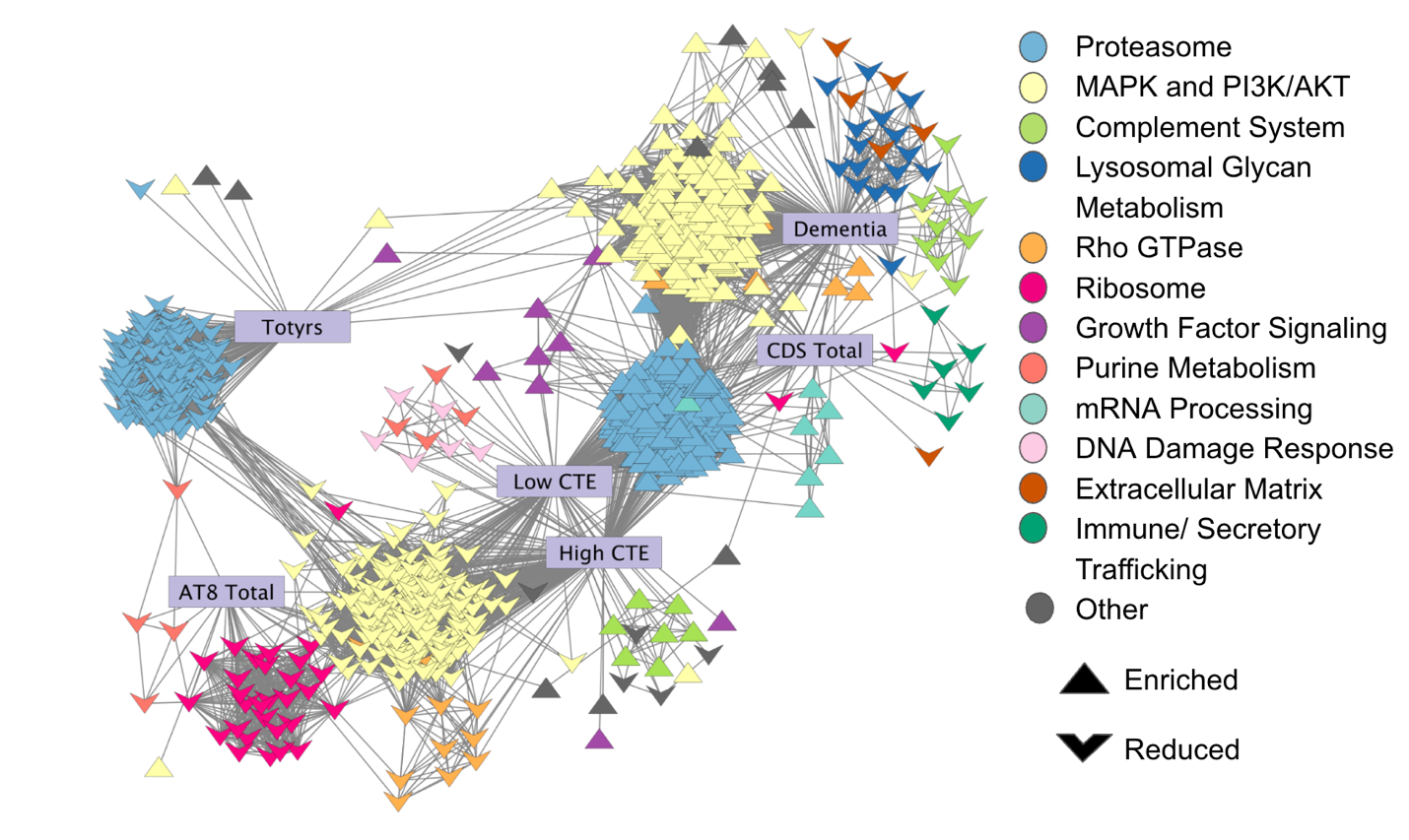

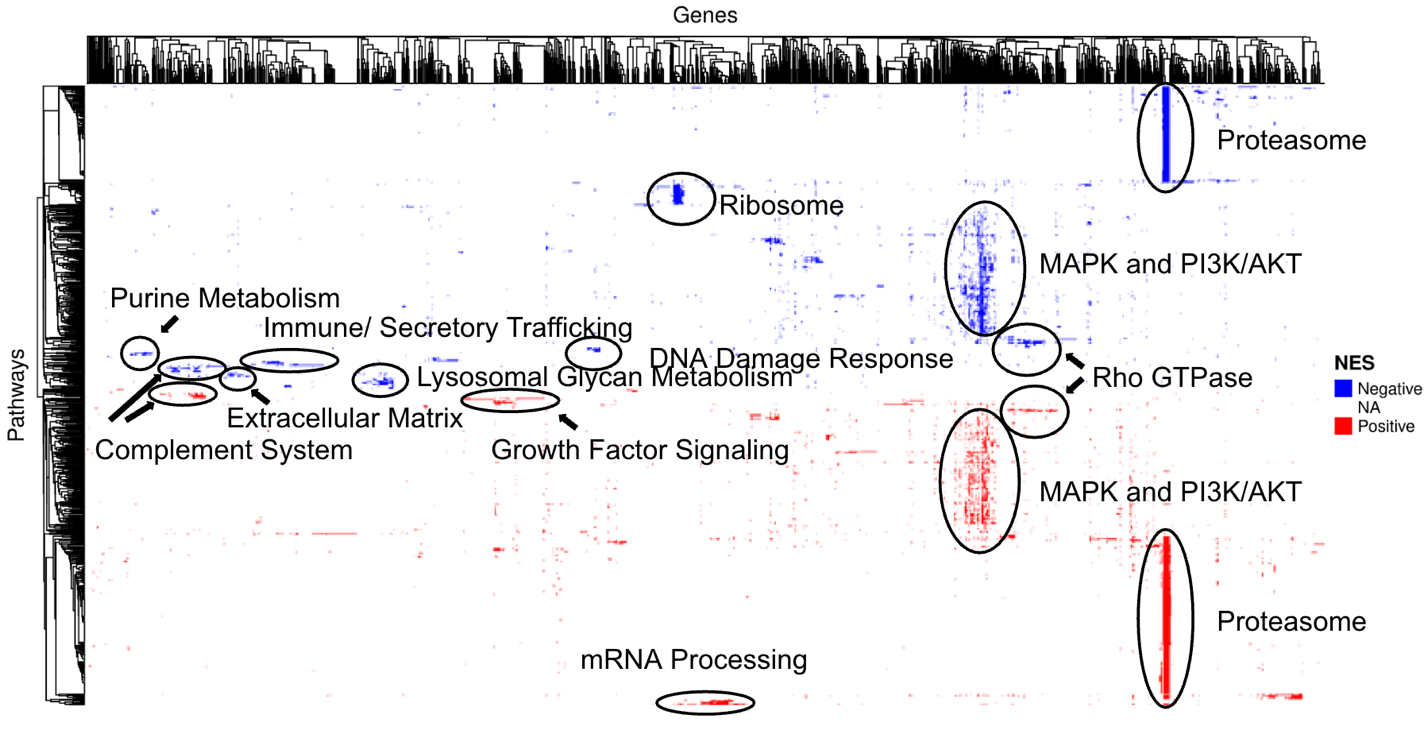
**Supplemental Figure 4. Network and Heatmap of all Hierarchically Clustered Significant Pathway Groups. (**Top) Network of all significant pathways across all six models in this study. Purple rectangles denote the six models; upwards facing triangles denote the enriched pathways, and downwards facing triangles denote reduced pathways. Significant pathways are connected if their cosine similarity score is greater than 0.2, and pathways are also clustered based on this value. Color coding of the pathways denotes which cluster of pathways they belong to, which is also based on the cosine similarity score hierarchical clustering. Groups needed to have at least five significant pathways. (Bottom) Heatmap of cosine similarity score hierarchical clustering of all significant pathways from all six models. Pathways are along the y-axis, and leading-edge genes are along the x-axis. Enriched pathways (positive NES) are red, reduced pathways (negative NES) are blue, and genes not in the leading edge remain white. Clusters are labeled with their group name.
